# Birth attendance in rural Bangladesh: practices and correlates

**DOI:** 10.1101/344788

**Authors:** Fozlul Korim

**Keywords:** delivery, childbirth, delivery attendance, determinants, rural area, Bangladesh

## Abstract

**Introduction:** The maternal mortality ratio (MMR) and neonatal mortality rate (NMR) are higher in the rural regions of Bangladesh compared to the urban areas or the national average. These two rates could be reduced by increasing use of skilled birth attendance in rural regions of this country. Although the majority of Bangladeshi population lives in rural areas, there has been a little investigation of the practices and determinants of delivery attendance in this region of Bangladesh. This study investigated the practices and determinants of attendance during child-births in rural Bangladesh.

**Methods:** Data were collected by the 2014 Bangladesh Demographic and Health Survey (BDHS 2014). After reporting the distribution of deliveries by types of attendance and distribution of selected factors, logistic regression was applied to calculate the crude and adjusted odds ratios (ORs) with 95% confidence intervals (CIs), and p-values.

**Results:** More than half of the deliveries (53.1%) were conducted by traditional attendants; community skilled attendants were present in only a small number of deliveries. The following factors were positively associated with deliveries by skilled attendants: 25-34 years age group of women (adjusted odds ratio (AOR): 1.4; 95% CI: 1.1-1.8), a higher education level of women (AOR: 2.9; 95% CI: 1.7-4.9), or their husbands (AOR: 2.4; 95% CI: 1.6-3.7), receiving antenatal care (AOR: 2.1; 95% CI: 1.6-2.7), and higher wealth quintiles (AOR of the richest wealth quintile vs the poorest: 3.5; 95% CI: 2.3-5.3). On the other hand, women having a higher parity (i.e., number of pregnancy, ≥2) led to a lower likelihood of delivery by skilled birth attendants. The proportion of deliveries attended by skilled attendants was significantly lower in the other six divisions compared to Khulna.

**Conclusions:** Socioeconomic factors should be considered to design future interventions to increase the proportion of deliveries attended by skilled delivery attendants. Awareness programs are required in rural areas to highlight the importance of skilled attendants. Further re-evaluation of the community skilled birth attendants program is required.

## Background

Bangladesh and other South Asian countries are experiencing an epidemiologic and demographic transition with a substantial reduction in maternal mortality ratio (MMR) and neonatal mortality rate (NMR) from1990 to 2015 (1,2). Reducing MMR and child mortality were targets of the Millennium Development Goals (MDGs). The Health Goals (Goal 3) of the United Nations’ (UN) Sustainable Development Goals (SDGs) also explicitly stated the targets to reduce the MMR and NMR to less than 70 per 100,000 live births and 12 per 1,000 live births, respectively (3). Despite having a substantial decline in reduction of under-five or maternal mortality, the neonatal mortality is almost static for the last couple of years in this country.

To reduce the MMR and NMR, the World Health Organization (WHO) highly recommends skilled birth attendants for deliveries. Skilled attendants are accredited health professionals who are trained to proficiently manage uncomplicated pregnancies, childbirth and the immediate postnatal period. They can also identify, manage and refer when there are any complications in women and newborns (4).

Bangladesh is a developing country of South Asia with an approximate area of 144,000 square-kilometers. The estimated population of this country is 160 million; nearly two-thirds of the people live in rural areas(5). Administratively, this country is divided into eight divisions. Similar to many other developing countries, Bangladesh does not have a national vital statistics system to find the health indicators in this country. Most the estimates come from the model based or from clustered surveys such as Bangladesh Demographic and Health Survey (BDHS) and Multiple Indicator Cluster Survey (MICS)(5). A large number of surveys revealed that there have been consistent inequalities between urban and rural areas in this country(5,6). For instance, the MMR and NMR in rural regions are substantially higher in rural region. On the other hand, the approach that has been found to effectively reduce these high MMR or NMR such as promotion of skilled attendants has been consistently lower in rural regions of this country(5). It is of particular importance to prioritize the indicators associated with health status of the rural people as the majority of people live in rural areas in Bangladesh (7) and MMR, NMR of this country are higher than most other countries (1,2).

In addition to the latest BDHS, it was one of the ubiquitous findings of the studies that investigated the practices and correlates of delivery attendance in Bangladesh found that delivery attendance in rural areas are particularly different compared to urban areas. Moreover, child birth is a complex process which had association with factors of multiple levels like individual, household, socioeconomic and community level; however, there have been a dearth of studies that specifically considered the practices and multi-level determinants of deliveries according to types of attendants across all rural regions of Bangladesh.

There have been a substantial investigation about the delivery attendants in this country; however, a lack of recent analysis about the delivery attendance particularly in rural regions indicates that newer studies are required. Earlier studies that examined the practices, patterns and determinants of delivery utilization in this country, found educational status of the women and their husbands, maternal occupation, wealth quintile, receiving antenatal care, a visit by family planning workers, transportation issues, and complications related to delivery are associated with care seeking behaviors for delivery(8–15). These factors could also influence child mortality or maternal knowledge level or contraceptive use or even many non-communicable diseases (16–27). The objectives of this study are to investigate the practices of delivery attendance in rural regions, to compare distribution according to types of delivery attendance, to identify the determinants that influence the delivery assistance, and to find the potential areas needed for intervention by the government to increase the proportion of deliveries by skilled attendants in rural areas.

## METHODS

### Survey Design

This study used data from the Bangladesh Demographic and Health Survey (2014 BDHS).

The secondary and cross-sectional data were collected as a part of seventh DHS. The details of the survey are available elsewhere (5).

The sample of the BDHS 2014 represented the demographics of women across the country. The survey used a sampling frame from the list of enumeration areas (EAs) of the 2011 Population and Housing Census of the People’s Republic of Bangladesh, provided by the Bangladesh Bureau of Statistics (BBS)(5). This survey is based on a two-stage stratified sample of households. In the first stage, 600 EAs were selected with probability proportional to the EA size, with 207 EAs in urban areas and 393 in rural areas. A complete household listing operation was then carried out in all the selected EAs to provide a sampling frame for the second-stage selection of households. In the second stage of sampling, a systematic sample of 30 households on average was selected per EA to provide statistically reliable estimates of key demographic and health variables for the country for urban and rural areas separately, and for each of the seven divisions. With this design, the survey selected 18,000 residential households, which were expected to result in completed interviews with about 18,000 ever-married women (5).

Initially, 17,989 households were selected, 17,565 of the households were occupied, and 17,300 (99%) of the households were then interviewed. A total of 18,245 ever-married women of reproductive age (15-49years) were identified in these households and 17,863 women were interviewed which achieved a response rate of 98%. There was similar response rate in rural and urban areas of the country.

In this analysis, we only included the deliveries that took place within the last three years preceding the survey in rural regions of the country by excluding the women who lived in urban regions during the survey. We examined their delivery attendance status along with the determinants of delivery attendance.

## STATISTICAL ANALYSES

In BDHS survey, women were interviewed about the attendants of their last childbirth. The ‘delivery attendant’ definition provided by the WHO was used for this analysis to define skilled and unskilled attendants; the BDHS also used the same definition. In this way, qualified doctors, nurses, midwives, paramedics, family welfare visitors (FWVs), and community skilled birth attendants (CSBAs) were included as skilled attendants. Then medical assistants, sub-assistant community medical officers (SACMOs), community health care providers (CHCPs), traditional birth attendants (TBAs), unqualified doctors, relatives, and neighbors were included as unskilled attendants. Table 1 shows list of all study variables.

**Table 1:**
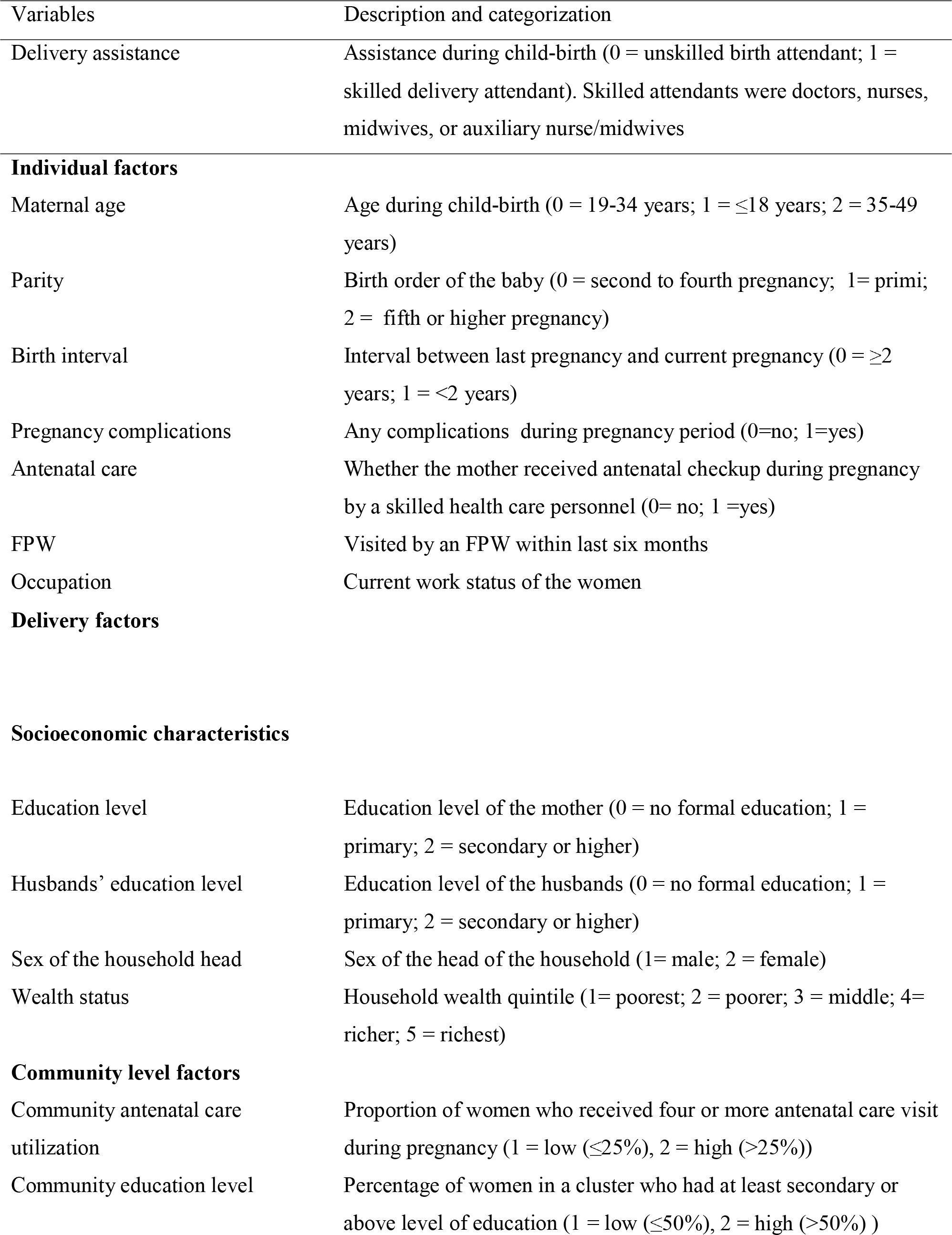
Study variables.

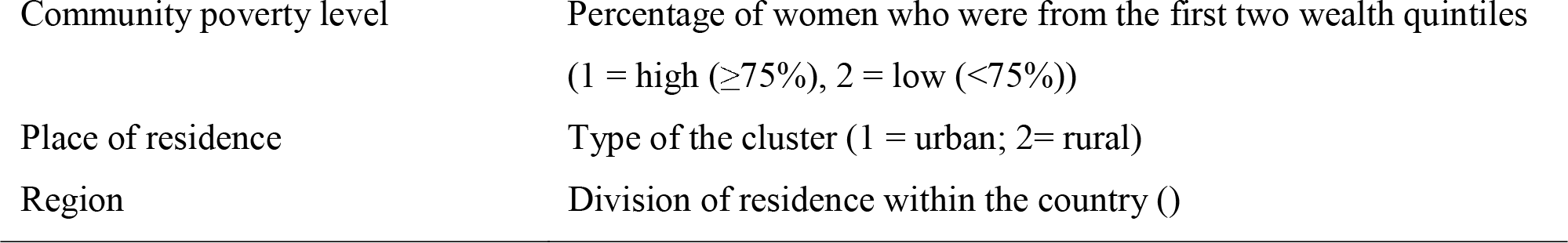

Stata 14.0 (Stata Corp, College Station, TX) was used to analyze data of this study (28). The ‘svy’ command was used to calculate the weighted frequency distribution. This command allowed adjustment for the two-stage stratified sampling of the survey design. Continuous (e.g., age) and discrete (e.g., parity) variables were converted into categorical variables. The wealth status was available a priori in the dataset; it was estimated by the principal component analysis (PCA) of the basic housing construction materials (i.e., materials used to construct the walls, roof, and floor of houses), sources of water, sanitation facilities, electricity, and household belongings.

At first, types of delivery attendance were reported (Figure 1). Then a contingency table was used to describe the background characteristics and to compare women according to these selected background characteristics. Simple logistic regression analyses were then applied to calculate crude (unadjusted) odds ratios (CORs). We considered the hierarchical data structure of the BDHS to construct multivariable model for this analysis; thus we performed a two-stage multi-level analysis. Covariates with a predetermined significance level (p<0.20) in bivariate analyses (i.e., simple logistic regression) were included for adjustment in multivariable analysis and the adjusted odds ratios (AORs) were reported (Table 2). To prevent residual confounding in multivariate models, the significance level of 0.2 has been considered adequate (29). Women who sought care from skilled attendants were compared to the women who did not seek care (skilled attendants =1, unskilled attendants = 0). Before including into the multivariate analysis, variable inflation factors (VIFs) were estimated to check multi-collinearity among the variables. Odds ratios (ORs) with 95% confidence intervals (CIs) were reported.

**Figure 1:**
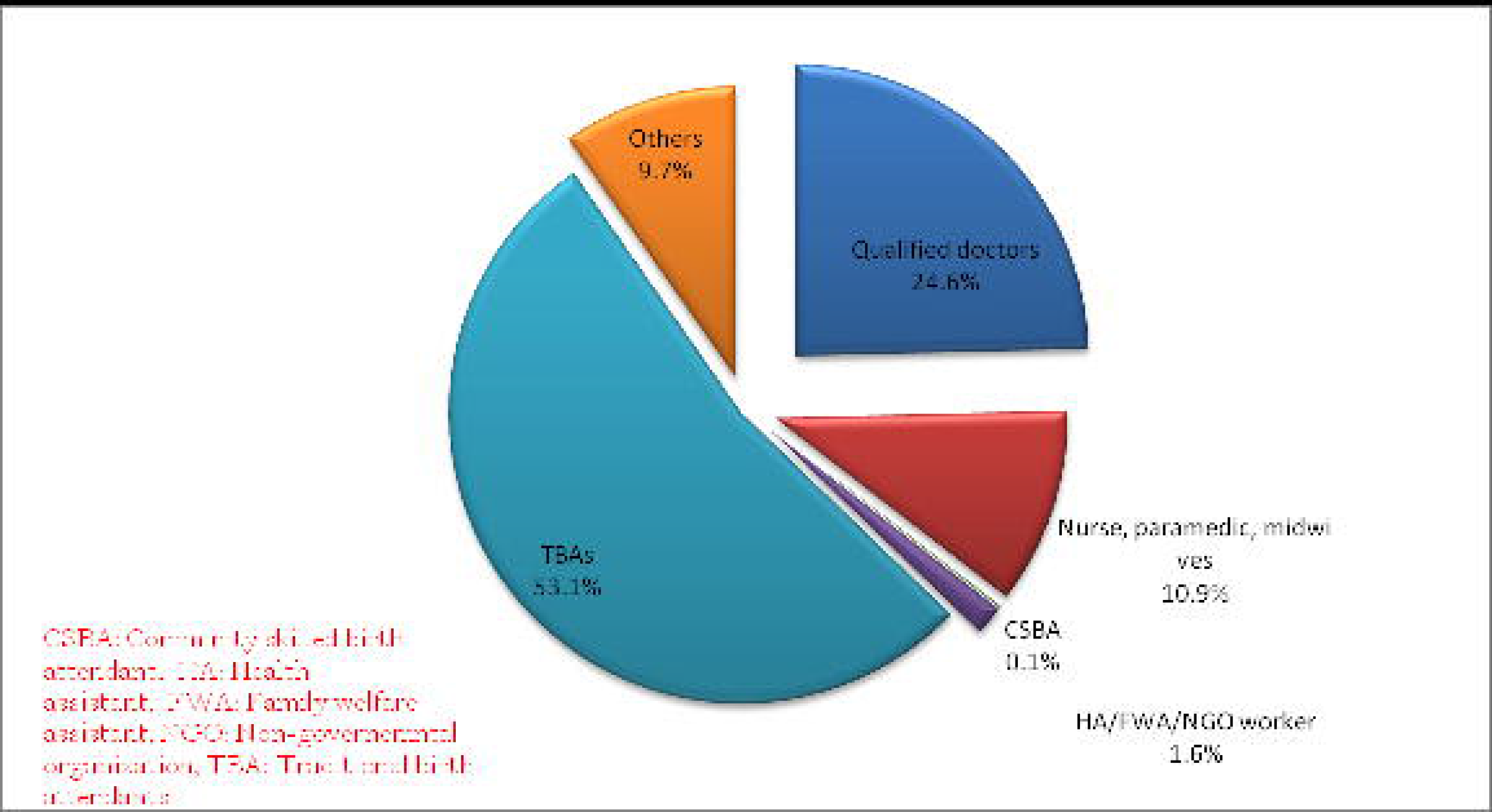
Distribution of deliveries by birth attendants.

**Table 2:** Weighted percentage distribution of deliveries by skilled birth attendants (SBAs), with selected background characteristics in rural Bangladesh

| Characteristics | Unskilled attendants | Skilled attendants | Total |
| --- | --- | --- | --- |
| **Current age** |  |  |  |
| 15-24 | 1,242 (62.6) | 742 (37.4) | 1,984 |
| 25-34 | 909 (64.8) | 494 (35.2) | 1,403 |
| 35-49 | 165 (73.8) | 59 (26.2) | 224 |
| **Parity** |  |  |  |
| 1 | 691 (53.6) | 598 (46.4) | 1,289 |
| ≥2 | 1,625 (70.0) | 697 (30.0) | 2,322 |
| **Birth interval (years)** |  |  |  |
| ≤2 | 311 (77.3) | 91 (22.7) | 402 |
| ≥3 | 2,005 (62.5) | 1,204 (37.5) | 3,209 |
| **Known pregnancy complications** |  |  |  |
| No | 1,674 (69.9) | 720 (30.1) | 2,394 |
| Yes | 642 (52.7) | 575 (47.3) | 1,217 |
| **Received antenatal care from a skilled provider** |  |  |  |
| No | 932 (83.3) | 186 (16.7) | 1,119 |
| Yes | 1,384 (55.5) | 1,109 (44.5) | 2,493 |
| **Received visit from a family planning worker** |  |  |  |
| No | 1,717 (64.7) | 918 (35.3) | 2,655 |
| Yes | 599 (62.6) | 357 (37.4) | 956 |
| **Occupation of the women** |  |  |  |
| Do not work for cash | 1,692 (62.3) | 1,023 (37.7) | 2,715 |
| Work for cash | 624 (69.6) | 272 (30.4) | 896 |
| **Education level of women** |  |  |  |
| No formal education | 476 (85.4) | 81 (14.6) | 557 |
| Primary | 800 (74.2) | 278 (25.8) | 1,078 |
| Secondary | 963 (56.3) | 746 (43.7) | 1,709 |
| College or above | 76 (28.6) | 190 (71.4) | 266 |
| **Education level of husbands of the women** |  |  |  |
| No formal education | 795 (80.4) | 193 (19.6) | 988 |
| Primary | 802 (69.7) | 348 (30.3) | 1,150 |
| Secondary | 598 (55.0) | 489 (45.0) | 1,087 |
| College or above | 120 (31.1) | 265 (68.9) | 385 |
| **Sex of the household head** |  |  |  |
| Male | 2,117 (64.1) | 1,186 (35.9) | 3,303 |
| Female | 199 (64.5) | 110 (35.5) | 309 |
| **Religion** |  |  |  |
| Islam | 2,155 (64.3) | 1,196 (35.7) | 3,351 |
| Others | 161 (61.8) | 99 (38.2) | 260 |
| **Division** |  |  |  |
| Barisal | 155 (74.1) | 54 (25.9) | 209 |
| Chittagong | 469 (60.5) | 306 (39.5) | 775 |
| Dhaka | 751 (65.1) | 403 (34.9) | 1,154 |
| Khulna | 135 (46.1) | 158 (53.9) | 293 |
| Rajshahi | 254 (65.0) | 137 (35.0) | 391 |
| Rangpur | 250 (64.2) | 140 (35.8) | 390 |
| Sylhet | 300 (75.6) | 97 (24.4) | 397 |

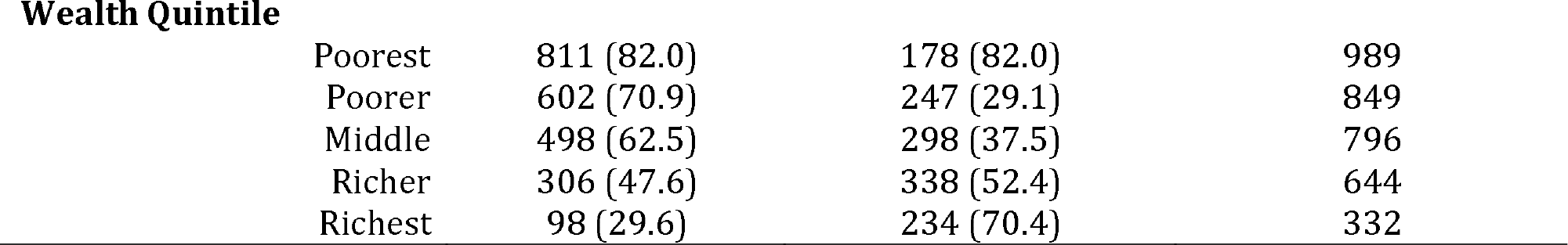

## RESULTS

A total of 3,611 weighted deliveries were included in our analysis. About 35.9% deliveries were attended by any types of medically trained provider (i.e., skilled attendants) in rural regions of this country (Figure 1). Among the skilled attendants, qualified doctors were the most common delivery attendants and attended 24.9% of the deliveries. Traditional birth attendants were the most common type (52.1%) of unskilled attendants. Community skilled birth attendants (CSBA) attended only 0.1% of the deliveries.

Table 2 provides selected background and socioeconomic characteristics of the respondents. The majority of the women were younger reproductive age group (15-24 years, 54.9%), followed by middle (25-34 years, 38.9%) and late (35-49 years, 6.2%) reproductive age groups. A total of 1,289 (35.7%) women were primi (i.e., pregnant for the first-time) and they received skilled delivery attendance in 46.4% of their deliveries which was higher than the non-primi mothers (30.4%). Approximately 69.0% women received antenatal care from a skilled provider during their pregnancy period. The women with a birth interval of three or more years received attendance from skilled attendants in 37.4% of the deliveries. The percentage of deliveries assisted by skilled attendants was higher among women or women with husbands who had any types of formal education compared to the women without any formal education. About 75.2% of the women were not involved with any type of occupation (i.e., do not earn money). The vast majority (92.8%) of the respondents were Muslims. More than fifty percent of the women were from poorest or poorer wealth quintile. The highest proportion of respondents were from Dhaka division (32.0%), followed by Chittagong (21.5%), Sylhet (11.0%), Rajshahi (10.8%), Rangpur (10.8%), Khulna (8.1%) and Barisal (5.9%), respectively. In comparison to other divisions, Khulna had the highest percentage of deliveries attended by skilled attendants, 53.9%. On the other hand, this proportion was less than one-fourth (24%) in Sylhet.

Table 3 shows the CORs, AORs, 95% CIs of the ORs and significance level (p-values) for the selected background and socioeconomic characteristics of the study participants. Age group was a significant predictor to be attended by skilled attendants in both adjusted and unadjusted levels. The women with an education level of college or above were three times more likely to use skilled birth attendants than the women without any formal education (AOR: 2.9; 95% CI: 1.7-4.9). Similar findings were observed for the education level of husbands. Women who were living in rural areas of Khulna division had a nearly three-fold higher likelihood of being delivered by SBAs than the women who lived in Barisal (AOR: 2.8, 95% CI: 1.8-4.4). The wealth status of women was also a significant predictor; the higher the wealth quintile, the higher the odds of being delivered by a medically trained provider except for the middle wealth quintile (AOR: 1.4, 95% CI: 0.9-2.2). There were no association between the likelihood of being delivered by skilled attendants and birth interval (AOR: 1.5, 95% CI: 0.9-2.5), occupation of the women (AOR: 0.9, 95% CI: 0.6-1.2), gender of the household head (AOR: 1.0; 95% CI: 0.7-1.4), and religion (COR: 1.1, 5% CI: 0.6-2.1).

**Table 3:** Logistic regression analyses showing the crude and adjusted odds ratios (with 95% confidence intervals) and significance level by delivery assistance in rural Bangladesh

| Characteristics | Crude odds ratio (95 % CI) | Adjusted odds ratio (95 % CI) |
| --- | --- | --- |
| **Current age (years)** |  |  |
| 15-24 | Ref | Ref |
| 25-34 | 0.9 [0.7,1.2] | 1.4* [1.1,1.8] |
| 35-49 | 0.6* [0.4,0.9] | 1.2 [0.8,2.1] |
| **Parity** |  |  |
| 1 | Ref | Ref |
| ≥2 | 0.5*** [0.4,0.7] | 0.6** [0.4,0.8] |
| **Birth interval (years)** |  |  |
| ≤2 | Ref | Ref |
| ≥3 | 2.1*** [1.4,3.0] | 1.5 [0.9,2.5] |
| **Known pregnancy complications** |  |  |
| No | Ref | Ref |
| Yes | 2.1*** [1.7,2.6] | 1.4** [1.1,1.7] |
| **Received ANC from a skilled provider** |  |  |
| No | Ref | Ref |
| Yes | 4.0*** [3.1,5.2] | 2.1*** [1.6,2.7] |
| **Received visit from a family planning worker** |  |  |
| No | Ref |  |
| Yes | 1.1 [0.7,1.6] |  |
| **Occupation of the women** |  |  |
| Do not work for cash | Ref | Ref |
| Work for cash | 0.7* [0.5,1.0] | 0.9 [0.6,1.2] |
| **Education level of women** |  |  |
| No formal education | Ref | Ref |
| Primary | 2.0*** [1.5,2.8] | 1.5* [1.1,2.1] |
| Secondary | 4.5*** [3.2,6.4] | 1.8** [1.2,2.5] |
| College or above | 14.6*** [9.1,23.4] | 2.9*** [1.7,4.9] |
| **Education level of husbands of the women** |  |  |
| No formal education | Ref | Ref |
| Primary | 1.8*** [1.4,2.3] | 1.2 [0.9,1.6] |
| Secondary | 3.4*** [2.4,4.7] | 1.5* [1.1,2.0] |
| College or above | 9.1*** [6.3,13.1] | 2.4*** [1.6,3.7] |
| **Sex of the household head** |  |  |
| Male | Ref |  |
| Female | 1.0 [0.7,1.4] |  |
| **Religion** |  |  |
| Islam | Ref |  |
| Others | 1.1 [0.6,2.1] |  |
| **Division** |  |  |
| Barisal | Ref | Ref |
| Chittagong | 1.9** [1.2,3.0] | 1.5* [1.0,2.3] |
| Dhaka | 1.5 [1.0,2.4] | 1.4 [0.9,2.0] |
| Khulna | 3.4*** [2.1,5.3] | 2.8*** [1.8,4.4] |
| Rajshahi | 1.5* [1.0,2.4] | 1.7* [1.1,2.5] |
| Rangpur | 1.6* [1.1,2.4] | 1.3 [0.9,2.0] |
| Sylhet | 0.9 [0.5,1.6] | 1.1 [0.7,1.6] |
| **Wealth Quintile** |  |  |

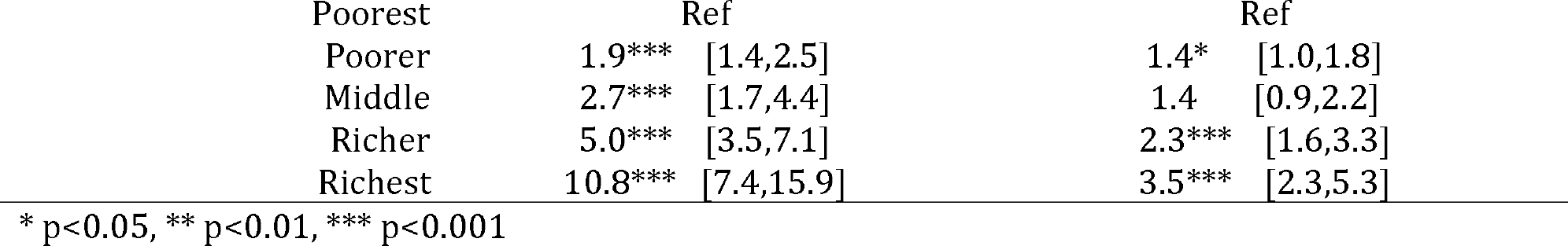

## DISCUSSION

The current study investigated a nationally representative dataset to investigate the practices and correlated of delivery attendance in rural regions of Bangladesh. Only one-third of the deliveries were attended by skilled attendants; this proportion is lower in comparison to most other countries, even lower than the urban areas or the national average of this country (5). TBAs attended more than half of the deliveries, and the proportion of births attended by CSBAs was very small. We found that age of the women, parity, education level of the women or their husbands, antenatal care by a skilled provider, division of residence and wealth status have significant associations with delivery attendance. We have reconfirmed the significance of several known correlates of delivery attendance for rural regions of Bangladesh.

The rural regions had lower utilization of facility delivery than the urban areas of the country which has resulted in the lower utilization of the skilled attendants during child-birth (5). We found that CSBAs assisted only 0.1% of the deliveries. The government of Bangladesh started CSBA training program with an aim to provide at least 2 CSBAs in each of the union of the country (30). A total of 7,000 CSBAs have been trained by the year 2012. Their lower utilization has been found by several studies from Bangladesh (9,31). A re-evaluation of the CSBA training program is required. However, it is also important to investigate the distribution of skilled attendants in the community as the large number of deliveries takes place at home.

This study found a negative association between higher parity (i.e., the number of times a woman became pregnant in her life) and delivery attendance. A positive association between pregnancy complications and first pregnancy was revealed previously (32,33). This finding could be linked to the utilization of skilled attendance or place of delivery with parity of the mother as our study also found a positive association between pregnancy complications and delivery attendance. Women with first pregnancy used more skilled attendants because of pregnancy complications which were difficult to manage for the traditional or unskilled attendants. This relationship between pregnancy complications and parity could help to explain their association with delivery attendance. Another explanation that has been put forward to explain the association between delivery attendance and parity is that when a woman is delivered by an unskilled attendant in her first pregnancy, then she is more likely to use unskilled attendants in her subsequent pregnancies (34).

Women who received antenatal check-up from a skilled provider were more likely to be delivered by skilled attendants than the women who were not checked by a skilled health care provider. Other studies also found a similar association (8,9,11,12). The increased likelihood of skilled attendance could have an association with receiving information and counseling from the health care providers regarding the importance of safe delivery attendance during that period. Antenatal care also helps to identify pregnancy complications (35). In addition, because of its effectiveness, antenatal counseling has been described as one of the main four pillars of the Safe Motherhood Initiative (36). However, the proportion of the women who received antenatal care was low (i.e., nearly one-fourth).

The finding of our study that education level of the women and their husbands had association with delivery attendance is supported by other studies (8,9,11). The educated couples might be more aware of the importance of skilled attendance during delivery. Furthermore, a higher education level enables men and women to have well-paying jobs and to obtain higher wealth status compared to people with a lower education level(37). This may be the other explanation of the association between education level and skilled delivery attendance in addition to the awareness; they were wealthy enough to bear the costs of skilled attendants as we have also found an association of delivery attendance with wealth status. In addition to this, education has been described as the strongest and most consistent predictor of the health status of humans (38).

As in other studies from Bangladesh, having a higher level of wealth (i.e., richer women) was positively associated with deliveries by skilled attendants (8,10,11). This association between delivery attendance and wealth status has been found in studies from other countries as well (34,39,40). The finding that no difference in the likelihood of skilled delivery attendance between women from middle wealth quintile and the lowest wealth quintile could be a spurious finding which has also been found elsewhere (34). Nearly 50% of the study women were from poorer or poorest wealth quintile which indicates that increasing the number of skilled delivery attendance is not achievable without improving the socioeconomic status of the people of the rural regions of the country. Though it is difficult to address and intervene to improve socio-economic status, a review of BRAC-ICDDR,B Joint Research Project Working Paper Series in Bangladesh revealed that microcredit programs could improve maternal and child health by improving the socio-economic conditions of the participants of the of the program (41). The government of the country could utilize that result and increase coverage of microcredit programs in rural areas to improve the socioeconomic conditions of the people to increase the presence of skilled attendants during deliveries, which would ultimately reduce the MMR and NMR of the country (42).

Women living in several divisions were less likely to be delivered by skilled attendants. Khulna was the only division with where more than fifty-percent of the deliveries were attended by skilled providers(16). In BDHS 2014, the proportion of women who received antenatal care was highest in Khulna in addition to the maximum percentage of facility delivery (5); these two factors have contributed to the skilled attendance during child-birth in rural regions of this division. The government should prioritize other areas to increase deliveries attended by skilled attendants(16,23).

The first and a major limitation of this study is that this study did not examine several factors which were revealed by other studies. For example, complications related to delivery, distance to the adjacent hospital, transportation issues to take the women to health facilities, presence of skilled attendants in the geographic area, and cost of skilled attendants which might be difficult to bear for the poor families were found to be associated with delivery attendance (8–11,13,14,43–45). Here we only analyzed data of survived women; we did not explore the determinants of the most affected group (i.e., women who died during child-birth), thus there may be potential survival bias in the data. Data of the BDHS survey were collected retrospectively and the data were cross-sectional; causality cannot be established because of uncertainty about temporal association.

The foremost strength of our study is that it is generalizable for the rural regions of the country; it covered the population from all divisions and the sample size of this study was large enough to adequately draw conclusions for the general population. The response rate of the BDHS survey was approximately 99%. Along with this high response rate, survey design and analytic strategy to include only women who delivered within the last three years enabled a minimal possibility of recall bias.

## CONCLUSIONS

This study showed practices and determinants of delivery attendance in rural regions of Bangladesh and found that skilled delivery attendance is still very low in rural Bangladesh. This study also found that socioeconomic factors mainly influence the delivery attendance in rural regions of Bangladesh in addition to other factors including antenatal care, parity, and pregnancy complications. Future interventions which would aim to increase skilled delivery attendance should consider these factors. More awareness programs are required in rural parts of the country regarding the significance of the skilled delivery attendance. It is also important to re-evaluate the CSBA training program for the country.

## Declarations

### Competing Interests

Noo competing interests to disclose.

### Competing Interests

The authors declare that they have no competing interests.

